# An Epitope-based Malaria Vaccine Targeting the Junctional Domain of Circumsporozoite Protein

**DOI:** 10.1101/2020.08.07.241802

**Authors:** Lucie Jelínková, Hugo Jhun, Allison Eaton, Nikolai Petrovsky, Fidel Zavala, Bryce Chackerian

## Abstract

A malaria vaccine that elicits long-lasting protection and is suitable for use in endemic areas remains urgently needed. Here, we assessed the immunogenicity and prophylactic efficacy of a vaccine targeting a recently described epitope on the major surface antigen on *Plasmodium falciparum* sporozoites, circumsporozoite protein (CSP). Using a virus-like particle (VLP)-based vaccine platform technology, we developed a vaccine that targets the junctional region between the N-terminal and central repeat domains of CSP. This region is recognized by monoclonal antibodies, including mAb CIS43, that have been shown to potently prevent liver invasion in animal models. We show that CIS43 VLPs elicit high titer and long-lived anti-CSP antibody responses in mice and non-human primates. Immunization with CIS43 VLPs confers partial protection from malaria infection in a mouse model, and both immunogenicity and protection were enhanced when mice were immunized with CIS43 VLPs in combination with adjuvants including delta inulin polysaccharide particles and TLR9 agonists. Passive transfer of serum from immunized macaques also inhibited parasite liver invasion in the mouse infection model. Our findings demonstrate that a Qß VLP-based vaccine targeting the CIS43 epitope combined with various adjuvants is highly immunogenic in mice and macaques, elicits long-lasting anti-CSP antibodies, and inhibits parasite infection in a mouse model. Thus, the CIS43 VLP vaccine is a promising pre-erythrocytic malaria vaccine candidate.

## Introduction

Malaria is a major global public health concern, causing 228 million infections and 405,000 deaths worldwide in 2018 ^1^. While malaria can be caused by several species of the parasitic organism *Plasmodium, P. falciparum* is responsible for causing a severe form of the disease with the highest morbidity and mortality, and is one of the leading causes of death in children under 5 years old ^1^. Infection is initiated when the female *Anopheles* mosquito injects sporozoites into the bloodstream of a human host. Sporozoites then rapidly transport to the liver where they transiently multiply within hepatocytes, producing merozoites. Merozoites then enter the blood stream where they invade erythrocytes, replicate further, and cause the symptoms and pathology of malaria ^2^.

Vaccines that target different stages of the malaria life cycle are under development ^2^. However, only vaccines which target the pre-erythrocytic stage have potential for providing sterilizing immunity ^2^. One of the primary targets of pre-erythrocytic vaccines is the major *P. falciparum* surface antigen circumsporozoite protein (*Pf*CSP) ^3,4^ CSP plays a critical role in facilitating parasite invasion of hepatocytes ^5^ and antibodies that target CSP can block infection ^6,7^. CSP consists of an immunodominant central repeat domain, which contains ∼38 copies of an NANP motif and up to four copies of an NVDP motif, flanked by N- and C-terminal domains ^8^. The sole approved malaria vaccine, RTS,S/AS01, is a recombinant protein-based vaccine developed in 1987 that is comprised of a large portion of CSP (consisting of 19 NANP repeats and the C-terminal domain) fused to the hepatitis B surface antigen (HBsAg) and co-expressed with wild-type HBsAg to form a particulate antigen which displays this CSP domain at low valency ^9^. RTS,S provides only modest protection (30-50%) from clinical malaria in endemic areas, and this protection has been shown to wane rapidly following immunization ^10–12^.

Most candidate CSP-targeted vaccines, including RTS,S, have focused on eliciting antibodies against the central repeat domain of the protein. Recently, however, potent inhibitory monoclonal antibodies (mAbs) which target a highly conserved epitope within the junctional region between the N-terminal domain and the central repeat region have been identified. MGG4 and CIS43 are mAbs that were isolated from human volunteers immunized with an experimental irradiated whole sporozoite vaccine (PfSPZ); these mAbs are highly effective at inhibiting liver invasion in mouse models of malaria infection ^13, 14^. These two mAbs bind to overlapping epitopes within the junctional domain, a region of CSP that is not included in the RTS,S vaccine ^4^. Interestingly, both antibodies also appear to be able to bind to the central NANP repeat domain, suggesting the possibility that this binding promiscuity contributes to the potent neutralizing activity of this class of antibodies ^15^. While the mechanism of antibody neutralization is not entirely clear, CIS43 has been shown to inhibit a proteolytic cleavage event in CSP that is critical for hepatocyte adhesion and, subsequently, invasion ^14^.

The discovery of a novel site of vulnerability within CSP has several clinical implications. First, neutralizing mAbs could be used as prophylactic treatment in humans traveling to malaria endemic areas. CIS43, for example, has recently entered Phase I trials for prophylactic prevention of clinical malaria in humans ^16^. Second, these data suggest that the junctional domain of CSP is a promising target for vaccine development. However, no effective vaccines targeting this region have been reported in the literature. Tan and colleagues immunized mice with a vaccine consisting of a 19 amino acid peptide encompassing the MGG4 epitope conjugated to keyhole limpet hemocyanin (KLH), but the antibodies elicited by this vaccine failed to block sporozoite invasion of hepatocytes ^13^. More broadly, there are significant challenges to developing an effective liver-stage vaccine targeting malaria sporozoites. Sporozoites can reach hepatocytes in less than an hour following infection ^17^, limiting the window of time in which immune responses can act. In addition, a single infected hepatocyte can seed the blood stage of the malaria lifecycle, meaning that effective immunity likely must be sterilizing ^2^. Thus, there is a high barrier for vaccine-mediated protection--an effective vaccine must elicit very high levels of circulating antibodies, and these antibodies must be long-lived ^18^.

As a potential solution, we investigated the effectiveness of an adjuvanted virus-like particle (VLP)-based vaccine targeting the CIS43 epitope. VLPs are non-infectious, self-assembling particles that are derived from viral structural proteins that can be used as standalone vaccines but also can be applied as platforms for vaccine development ^20, 21^. VLP-based vaccine design exploits the intrinsic ability of viral structural proteins to self-assemble into highly immunogenic, multivalent particles. These multivalent structures are effective at stimulating strong antibody responses by promoting B cell receptor crosslinking, leading to robust and long-lasting antibody responses against diverse target antigens ^19–21^. In addition, adjuvants play a key role in maximizing the ability of vaccines to induce high titer, high avidity antibodies that are durable over time. In the past most human malaria vaccine candidates have incorporated either aluminium salts (alum) or oil emulsion adjuvants such as Montanide. However, the former have, by and large, proved insufficiently immunogenic^22^, while the latter have suffered from high reactogenicity^23^, raising safety concerns. In recent years a variety of new adjuvants have become available, raising the possibility to use these to improve malaria vaccine efficacy without compromising safety. Advax adjuvants are proprietary adjuvant formulations produced by Vaxine Pty Ltd, Australia, that are based on particles including those made from inulin, a plant-based polysaccharide ^24,25^. These particulate adjuvants have been shown to be potent enhancers of both cellular and humoral immunity. These particles are co-formulated with a range of other immune modulators such as TLR9 active CpG oligonucleotides to achieve synergistic adjuvant effects and help steer the immune response in any desired direction.

Here, we describe the development and characterization of a bacteriophage VLP-based vaccine targeting the CIS43 epitope. CIS43 VLPs elicit high-titer antibody responses against *P. falciparum* CSP in both mice and macaques, particularly in combination with Advax adjuvants. These antibody responses are highly durable, and vaccination inhibits malaria invasion of the liver in a mouse model.

## Results

### Construction and antigenicity of CIS43 VLPs

The CIS43 mAb epitope was mapped to a 15 amino acid peptide at the N-terminus of the repeat domain of CSP, spanning CSP amino acids 101-115 ^14^. To assess whether a vaccine targeting this epitope could elicit antibodies with CIS43-like activity, we engineered RNA bacteriophage VLPs to display the CIS43 epitope at high valency. A peptide representing CSP_101-115_ was synthesized to contain a short gly-gly-gly-cys linker sequence and then conjugated to the surface lysines on Qß bacteriophage VLPs using a bifunctional crosslinker (shown schematically in Fig. 1A) to produce CIS43 VLPs. Conjugation efficiency was measured by SDS-PAGE analysis.

**Fig. 1.**
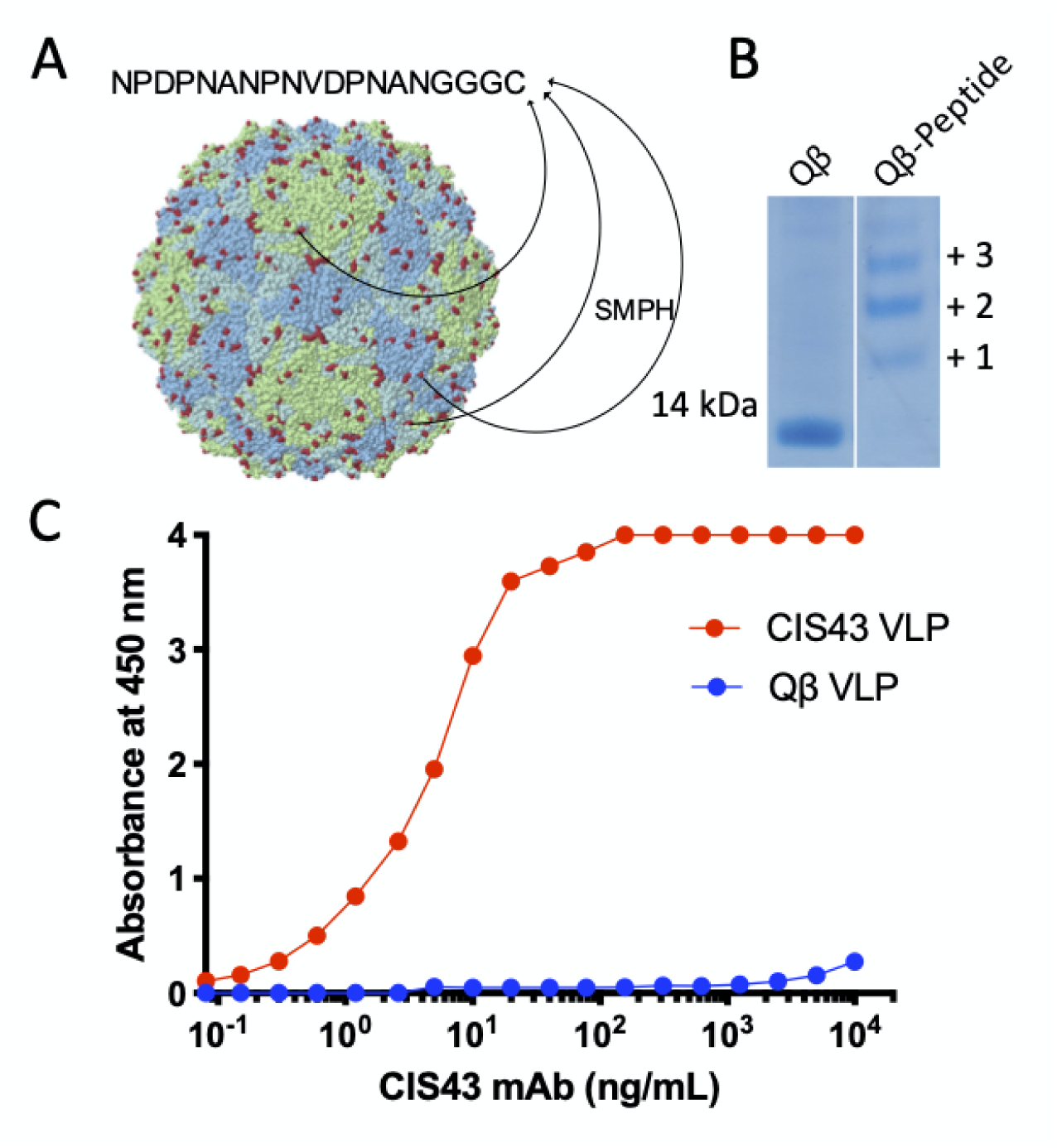
Characterization of CIS43 VLPs. (**A**) Schematic representation of CIS43 VLP conjugation. A 15-amino acid peptide representing the CIS43 mAb epitope was synthesized to include a (glycine)_3_-cysteine linker and conjugated to surface-exposed lysine residues (shown in red) on the coat protein of Qß bacteriophage VLPs using the bifunctional crosslinker SMPH. (**B**) SDS-PAGE analysis of unconjugated (left lane) or CIS43 peptide conjugated (right lane) Qß VLPs. The ladder of bands in the CIS43 VLP lane reflect individual copies of coat protein modified with 1, 2, or more copies of the CIS43 peptide. (**C**) Binding of the CIS43 mAb to CIS43 VLPs (red) or wild-type (unmodified) Qß VLPs (blue) as measured by ELISA.

Successful peptide conjugation is indicated by an increase in the molecular weight of Qß coat protein subunits, reflecting conjugation of one or more peptides to Qß coat protein (Fig. 1B, righ lane). More than half of all coat protein bound two or more copies of peptide, suggesting that the particles are decorated with the peptide in a dense and multivalent fashion. We estimate that an average of 360 copies of the peptide were conjugated to each Qß VLP.

To assess the antigenicity of the CIS43 VLPs, we measured the binding of CIS43 mAb to CIS43 VLPs by ELISA. As shown in Fig. 1C, CIS43 VLPs were robustly recognized by the CIS43 mAb. This suggests that the CIS43 epitope peptide is displayed on the Qß particles in a manner that emulates its antigenic conformation on CSP.

### CIS43 VLPs induce high titer antibody responses against CSP

We have previously shown that multivalent display of peptides on VLPs can stimulate high titer antibody responses. To assess the immunogenicity of the CIS43 VLPs in mice, Balb/c mice were intramuscularly immunized with CIS43 VLPs or, as a negative control, wild type (WT) Qß VLPs and boosted at 3 and 7 weeks (Fig. 2A). Three weeks following the final boost, antibody responses against the CIS43 peptide and full-length recombinant *P. falciparum* CSP were measured by end-point dilution ELISA. Whereas negative control sera collected from mice immunized with WT Qß VLPs did not show reactivity to the CIS43 peptide (not shown) or full-length CSP, CIS43 VLPs elicited strong anti-peptide (Fig. 2B and Supplementary Fig. 1) and anti-CSP antibody responses (Fig. 2B). The antibody titer to CSP was slightly higher than the anti-CIS43 peptide titer, possibly because a portion of the CIS43 epitope shares homology to the CSP repeat domain ^14^. Similar antibody titers were observed in C57BL/6 mice immunized with CIS43 VLPs (data not shown). To interrogate the binding promiscuity of elicited anti-CSP antibodies from immunized Balb/c mice, we tested the ability of sera to bind to several peptides representing epitopes within and flanking the junctional domain of CSP (shown in Supplementary Fig. 1) by ELISA (these epitopes are described in references ^9, 13, 14, 23^). While antisera bound most strongly to the peptide corresponding to the CIS43 epitope, we also detected cross-reactive binding to several other CSP-derived peptides, particularly those peptides which contained one or more of the NANP motifs found in the CSP central repeat domain (Supplementary Fig. 2).

**Fig. 2.**
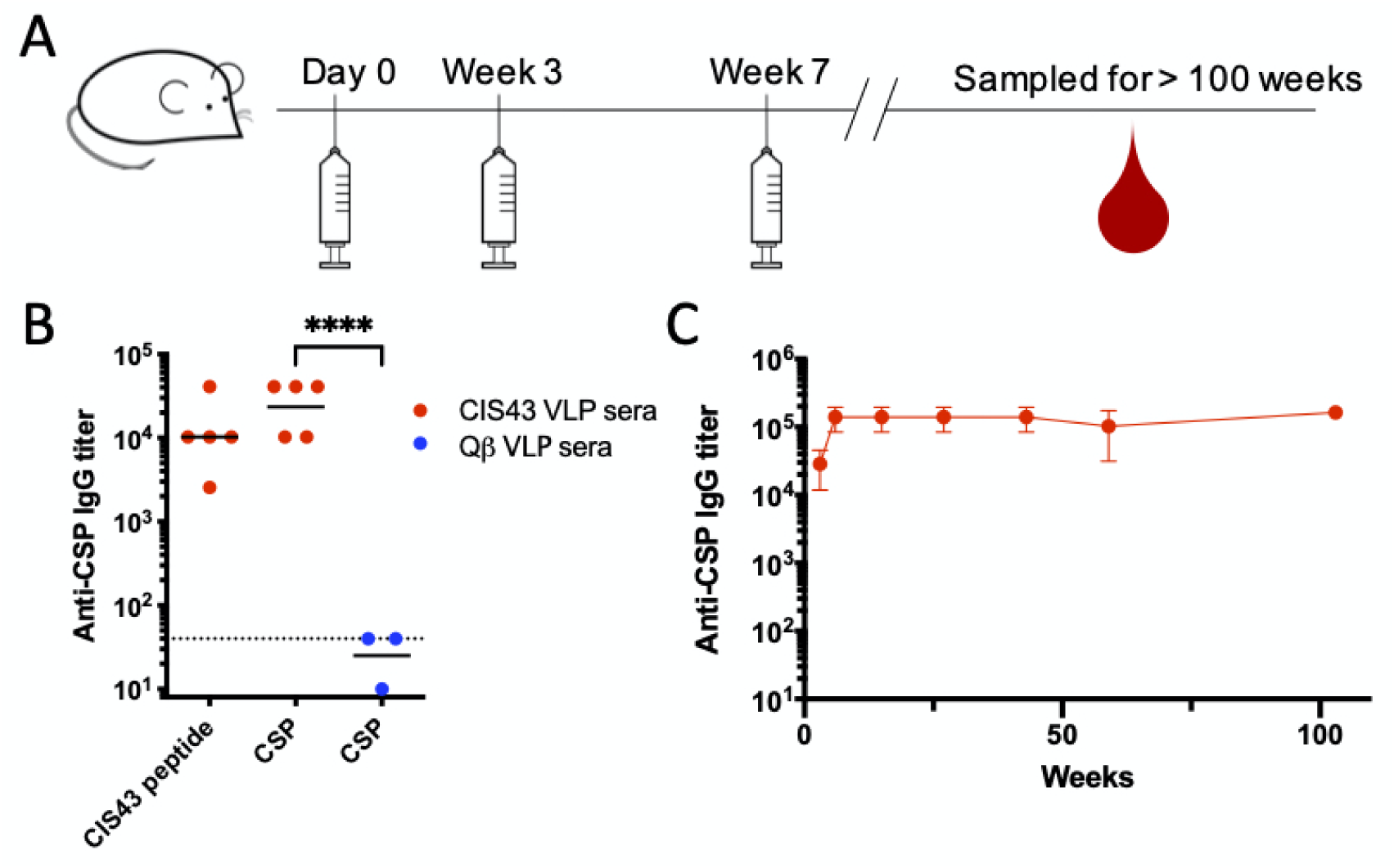
CIS43 VLPs elicit high titer and long-lasting antibody responses against CSP. (**A**) CIS43 VLP immunization scheme. (**B**) IgG titers against CIS43 peptide or CSP were determined using sera collected 6 weeks after the initial vaccination. Balb/c mice were immunized with 5 µg of CIS43 VLPs (n = 5; red symbols) or 5 µg of WT Qß VLPs (n = 3; blue symbols). Each data point represents an individual mouse and lines represent the geometric mean titers of each group. The dashed line indicates the limit of detection of the ELISA. Statistical significance between groups was determined by t-test (****; *p* < 0.0001) (**C**) Geometric mean IgG titers were determined longitudinally for over 100 weeks after the initial vaccination. Note that three of the mice in the CIS43 VLP vaccinated group were sacrificed or died (at weeks 56, 60, & 60) due to health complications unrelated to vaccination and prior to the completion of the study.

We and others have previously shown that vaccination with VLPs can elicit differentiation of long-lived plasma cells (LLPCs), resulting in stable, long-lasting antibody levels ^27–29^. It is likely that long-lived circulating antibodies will be required for sustained protection from malaria infection in endemic areas, where reinfection is common. We examined the longevity of the antibody response to CIS43 VLPs in mice. Serum was collected regularly for two years after the third immunization and anti-CSP IgG titers were determined by ELISA. Remarkably, anti-CSP IgG titers were stable over this period, nearly spanning the lifetime of the mice (Fig. 2C). Together, these experiments illustrate that CIS43 VLPs elicit robust and long-lived antibody responses.

### Immunization with CIS43 VLPs protects mice from intravenous challenge with Plasmodium

To examine whether immunization with CIS43 VLPs confers protection from infection, we took advantage of a well-characterized mouse infection model for testing CSP-targeted vaccines. In this model, mice are challenged with transgenic *P. berghei* (*Pb*) that have been engineered to express full-length *Pf*CSP (in place of *Pb*CSP) and a luciferase reporter (*Pb-Pf*CSP-*Luc*) ^30^. 42 hours after infection, liver-stage parasite burden can be quantitated by measuring luciferase levels. Mice were immunized three times with CIS43 VLPs or, as a negative control, WT Qß VLPs, and then were challenged by intravenous injection of 1000 *Pb-Pf*CSP*-Luc* sporozoites four weeks following the final vaccine boost (Fig. 3A). Liver luciferase levels were compared to unimmunized (naïve) mice. Mice immunized with CIS43 VLPs had a significantly reduced liver-stage parasite burden compared to naive mice or controls vaccinated with WT Qß VLPs (Fig. 3B and 3C). No statistical difference was observed between naïve infected mice and mice immunized with WT Qß VLPs. These data suggest that immunization with unadjuvanted CIS43 VLPs provides partial (∼68%) inhibition of parasite liver invasion and protection from malaria infection in a rigorous challenge mouse model.

**Fig. 3.**
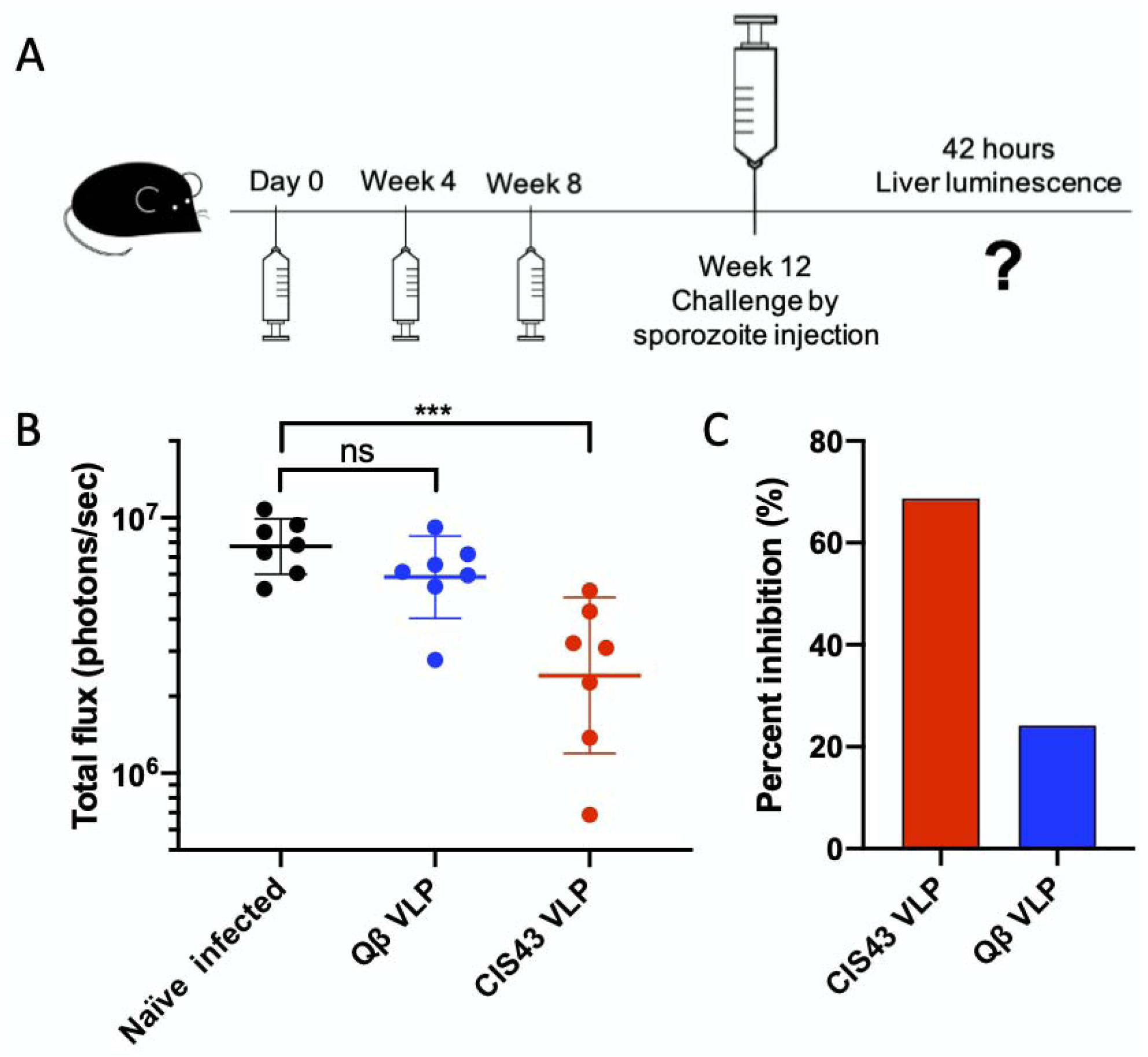
Immunization with CIS43 VLPs without exogenous adjuvant confers partial protection from *Plasmodium* infection. (**A**) Immunization and challenge scheme. (**B**) Parasite liver load (as measured by luminescence) in vaccinated or naïve mice following challenge by sporozoite injection. Horizontal lines represent the geometric mean luminescence of each group, error bars represent standard deviation of the geometric mean. Baseline luminescence was measured at 10^5^ photons/sec. Groups were compared statistically using one-way ANOVA followed by Dunnett’s multiple comparison test. ns; not significant, ***; p<0.001. (**C**) Percent inhibition of liver infection, as calculated from luminescence data. Inhibition of infection is calculated relative to the geometric mean signal in the naïve, challenged group of mice.

### Advax adjuvants increase the immunogenicity of CIS43 VLPs

Immunization with CIS43 VLPs provided partial protection from *in vivo* challenge. Previous research showed that the protection conferred by passive antibody transfer of CIS43 mAb is dose dependent ^14^, suggesting that higher antibody levels could potentially provide stronger protection. As a strategy to increase antibody titers elicited by CIS43 VLPs, we tested the compatibility of CIS43 VLPs with Advax adjuvants. Advax adjuvants contain delta inulin polysaccharides that have been shown to enhance both the cellular and humoral immune responses to a variety of antigens and can be combined with other immunostimulatory agents, such as CpG oligonucleotides ^28, 29^. Mice were immunized three times with CIS43 VLPs in combination with five different Advax formulations (Advax 1-5), CpG oligonucleotides alone, or without exogenous adjuvant, and then antibody levels were determined by ELISA. As shown in Supplementary Fig. 3, Advax adjuvants, including those containing CpG, but not CpG by itself, increased anti-CSP antibody titers relative to unadjuvanted CIS43 VLPs. All of the vaccine formulations induced statistically similar IgG1/IgG2a ratios (Supplementary Fig. 4).

Amongst the adjuvants that were tested, Advax-3 and Advax-4 had the greatest boosting effect on anti-CSP titers (Supplementary Fig. 3). To more carefully measure the impact of adjuvants, we quantitated anti-CSP antibody concentrations in vaccinated mice by linear regression analysis using a standard curve generated with the mouse anti-CSP mAb 2A10 ^32^. After two immunizations, mice immunized with unadjuvanted CIS43 VLPs generated a mean anti-CSP IgG level of 44 µg/mL. Formulation of CIS43 VLPs with Advax-3 or Advax-4 significantly boosted anti-CSP IgG concentrations by 5.5-fold and 2.3-fold, respectively (Fig. 4A).

**Fig. 4.**
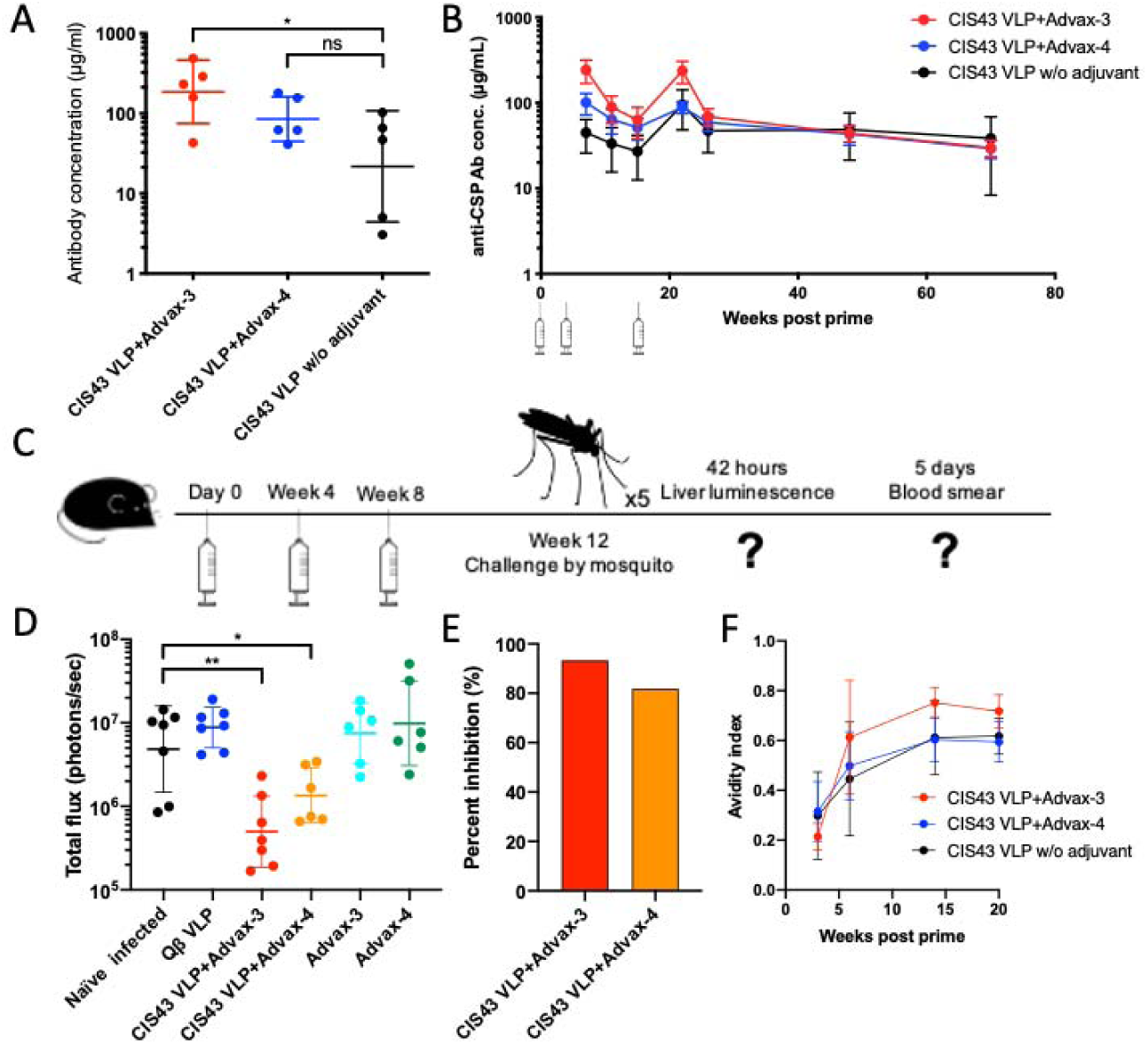
Advax adjuvants enhance both CIS43 VLP immunogenicity and protection from malaria infection. (**A**) Anti-CSP antibody concentrations in Balb/c mice immunized two times with CIS43 VLPs with or without Advax-3 or Advax-4 (n = 5/group). ns; not significant, *; p<0.05. (**B**) Kinetics of anti-CSP antibody concentrations in Balb/c mice immunized three times with CIS43 VLPs with or without adjuvant. Mice were immunized at week 0, 4, and 18, and anti-CSP antibody concentrations were measured at various timepoints following immunization. (**C**) Immunization and challenge scheme. Groups of mice (n=6 or 7/group) were immunized three times and then challenged with five *Pb-Pf*CSP*-Luc* infected mosquitos. Liver luminescence was evaluated 42 hours following mosquito challenge. (**D**) Parasite liver load (as measured by luminescence) in CIS43 VLP vaccinated or control mice following mosquito challenge. Mann-Whitney test was used to statistically compare each group to the unimmunized (naïve) group. *; p<0.05, **; p<0.01. (**E**) Percent inhibition of liver infection, as calculated from luminescence data. Inhibition of infection is calculated relative to the geometric mean signal in the negative control groups of mice. (**F**) Avidity index (AI) of anti-CSP antibodies elicited following immunization with CIS43 VLPs with or without adjuvant.

To evaluate the longevity of the response, we followed anti-CSP IgG responses over time. As seen in Fig. 4B, antibody responses induced by unadjuvanted CIS43 VLPs were stable over time. Advax adjuvants, particularly Advax-3, provided a boost in antibody responses that declined following the second immunization, suggesting that some of the antibody responses were secreted by short-lived plasma cells. Antibody titers rebounded to peak concentrations after a second boost, 18 weeks after the primary immunization. As we showed previously, CIS43 VLPs elicited anti-CSP IgG concentrations which were remarkably stable following the initial decline after the second boost (Fig. 4B).

### Vaccination with CIS43 VLPs combined with Advax-3 or Advax-4 more strongly inhibits Plasmodium infection in mice

To evaluate the protection elicited by CIS43 VLPs adjuvanted with Advax-3 or Advax-4, C57BL/6 mice (n = 6-7/group) were vaccinated with CIS43 VLPs with adjuvant, or with adjuvant alone, and then were challenged with *Pb-Pf*CSP*-Luc* sporozoites. In this experiment, mice were challenged with live parasite-carrying mosquitoes in order to more closely recapitulate the conditions of natural infection (Fig. 4C). Mice immunized with CIS43 VLPs adjuvanted with Advax-3 or Advax-4 showed significantly lower parasite liver loads compared to the naïve group, mice immunized with either adjuvant alone, or wild-type Qß VLPs (Fig. 4D). Animals immunized with CIS43 VLPs adjuvanted with Advax-3 or Advax-4 showed a 90% and a 72% reduction in liver parasite loads compared to naïve mice, respectively. When we compared liver loads in these two groups with an aggregate of all of the negative control groups, we measured an 93% (CIS43 VLPs with Advax-3) and a 82% (CIS43 VLPs with Advax-4) reduction (Fig. 4E). Despite these reductions in liver load, all of the mice in this study developed parasitemia by day 5 post-infection, as evaluated by blood smears. These results demonstrate that the CIS43 VLP vaccine is capable of stimulating high antibody responses that confers a robust, although not sterilizing, level of protection from liver infection in a mouse model.

It was previously reported in clinical trials of RTS,S/AS01 that increased antibody avidity, along with antibody concentration, correlated with increased protection from clinical malaria ^18,33^. To understand how avidity of CIS43 VLP-elicited antibody changes over time and in response to added adjuvant, we evaluated the avidity index (AI) of antibodies against CSP using a chaotrope-based avidity assay. The mean AI of each group increased after each immunization, culminating in a high mean AI value (>0.5) in all of the groups following the third immunization (Fig. 4F). Amongst the three vaccine groups we tested, Advax-3 adjuvanted CIS43 VLPs elicited anti-CSP antibodies with the highest avidity (Fig. 4F).

### CIS43 VLPs elicit anti-CSP antibody responses in non-human primates and these antibodies can protect mice from malaria infection upon passive transfer

CIS43 VLP immunogenicity in non-human primates was evaluated by immunizing groups (n = 3) of cynomolgus monkeys with unadjuvanted CIS43 VLPs or CIS43 VLPs adjuvanted with Advax-3. All of the macaques developed anti-CSP antibodies (Fig. 5A), but there was some heterogeneity in the responses, possibly due to the diverse ages and sizes of the animals in the study (Supplementary Table 1). To investigate whether serum from CIS43 VLP-immunized cynomolgus monkeys could mediate protection from infection, we performed a passive transfer of monkey sera into naïve C57BL/6 mice and then challenged the mice with *Pb-Pf*CSP*-Luc* by mosquito infection. Macaque sera was obtained 2 weeks after the second immunization, pooled by immunization group, and then was used to passively immunize mice. Following intravenous administration of this pooled macaque sera, mice were rested for two hours and then subjected to mosquito challenge. As before, parasite liver load was evaluated by liver luminescence. We measured a statistically significant decrease in parasite liver burden in mice passively immunized with pooled sera from macaques immunized with unadjuvanted CIS43 VLP group and Advax-3-adjuvanted CIS43 VLPs, but not using sera from macaques immunized with control WT Qß VLPs. Sera from monkeys immunized with unadjuvanted CIS43 VLPs reduced liver parasitemia by 80%, and sera from Advax-3-adjuvanted CIS43 VLPs reduced liver parasitemia by 83%. Thus, these data show that non-human primates immunized with CIS43 VLPs elicit antibody responses that are capable of blocking hepatocyte invasion.

**Fig. 5.**
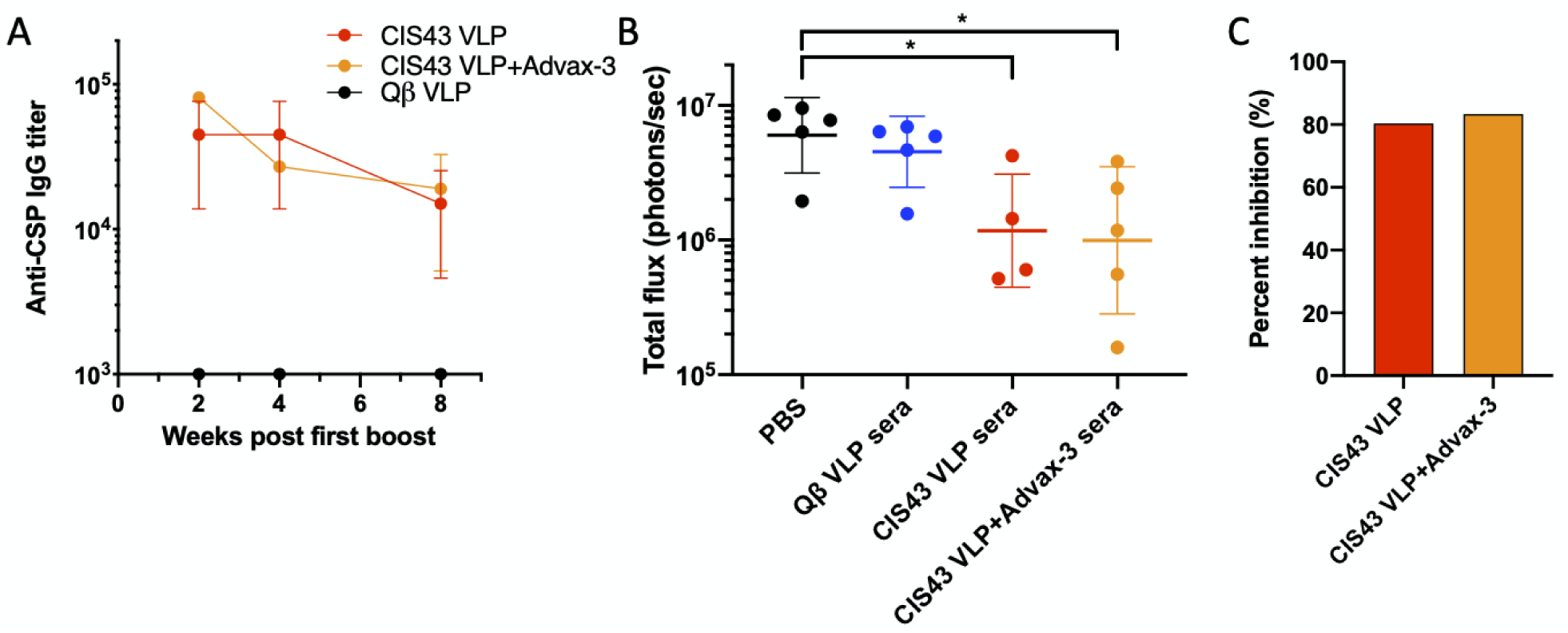
Cynomolgus monkeys immunized with CIS43 VLPs produce protective anti-CSP antibodies. (**A**) IgG concentrations in cynomolgus monkeys immunized at weeks 0 and 4 with CIS43 VLPs with or without Advax-3 adjuvant (n = 3/group). Qß VLPs were used as a negative control. (**B**) Parasite liver load (as measured by luminescence) in mice (n = 4-5/group) that received an intravenous injection of 500 µl of sera from immunized cynomolgus monkeys. Two hours following serum transfer, mice were challenged with five *Pb-Pf*CSP*-Luc* infected mosquitoes. Mann-Whitney test was used to statistically compare each group to the group that received an intravenous injection with PBS. (C) Percent inhibition of liver infection, as calculated from luminescence data. Inhibition of infection is calculated relative to the geometric mean signal in the groups of mice treated with PBS.

## Discussion

Development of an effective vaccine for malaria has been complicated by a number of features of the parasite, including its complex lifecycle, antigenic variability of its surface proteins, and the fact that immunity to natural infection largely does not confer protection from reinfection. One of the goals of an epitope-based malaria vaccine is to direct immune responses to target vulnerable, conserved domains of the pathogen. Here, we evaluated the efficacy of a novel vaccine targeting a highly conserved epitope ^14, 32^ within the junctional domain of *Plasmodium falciparum* CSP that is recognized by a potent inhibitory monoclonal antibody, CIS43. We showed that a VLP-based vaccine that displays the CIS43 epitope elicited high-titer antibodies against CSP in both mice and non-human primates and that vaccination inhibited parasite invasion in a mouse *Plasmodium* challenge model. Passive immunization of mice with CIS43-immunized macaque sera also inhibited infection, indicating that antibodies mediate these prophylactic effects. The mechanism(s) by which antibodies induced by CIS43 VLPs provide protection is unclear, but it has been shown that the CIS43 mAb can partially inhibit proteolytic cleavage of CSP ^14^, which is critical for invasion of hepatocytes ^35^. CIS43, and other mAbs that target the junctional domain of CSP ^15^, also cross-reacts with the central repeat domain of CSP. Antibodies that bind to the central repeat domain CSP can block infection by inhibiting sporozoite motility ^18, 34^, inducing parasite cytotoxicity ^37^ and/or by enhancing immune clearance ^35, 36^. We showed that antibodies induced by CIS43 VLPs bound most strongly to the CIS43 epitope, but they also cross-reacted with other CSP epitopes in the junctional domain and the central repeat domain, raising the possibility that this binding promiscuity may contribute to their inhibitory activity. Producing a vaccine that specifically targets the R1 cleavage site in the junctional domain could potentially dissect the relative contributions of how antibody binding to these two regions contributes to protection.

The ability to elicit durable antibody responses through the induction of long-lived antibody secreting plasma cells is likely to be particularly important for providing protection from malaria infection. Liver infection occurs within a few minutes or hours of parasite exposure and once *Plasmodium* invades hepatocytes it is not accessible to anti-sporozoite antibodies ^40^. Thus, circulating antibodies, but not reactivated memory B cells, are likely to be a critical mediator of effective immunity to malaria sporozoites. Indeed, one of the primary shortcomings of RTS,S is that antibody levels decrease rapidly following immunization ^41^. Here, we showed that CIS43 VLPs not only elicited high titer antibody responses against CSP, but that these antibodies were extremely durable. Over the span of a nearly two-year period following vaccination, mouse anti-CSP antibody titers did not decline. These data add to a growing body of literature demonstrating that the ability to induce long-lived plasma cells is a feature of vaccine antigens, such as VLPs, with highly multivalent structures ^27–29^. Our study provides evidence that use of a multivalent vaccine technology can effectively elicit durable antibody responses against CSP.

There have been several other efforts to develop multivalent vaccines targeting CSP. R21 is a RTS,S-like vaccine in which the central repeat domain of CSP is displayed at much higher density on recombinant hepatitis B surface antigen particles than RTS,S. R21 elicits potent immune responses against CSP ^42^ and is currently in Phase 1/2a clinical trials. Several groups have applied a similar approach using bacteriophage VLP-based vaccines to display full-length CSP at high valency and have shown that these vaccines are immunogenic in mice ^43^ and can confer protection following transgenic parasite challenge ^44^. Whitacre and colleagues developed epitope-targeted vaccines that employ woodchuck hepatitis virus core antigen (WHcAg) VLPs to display B and T cell epitopes from repeat and non-repeat domains of CSP ^9^. VLPs displaying repeat, but not non-repeat, epitopes were capable of eliciting sterilizing immunity that prevented blood stage infection in a mouse model. These latter challenge studies utilized a transgenic *Pb* CSP that only contained the repeat region of *Pf*CSP (unlike our studies in which transgenic *Pb* carried full-length *Pf*CSP). Thus, it is difficult to compare the relative protection provided by WHcAg-based VLPs versus the CIS43 VLPs described in this study. Nevertheless, a common finding is that VLP display of CSP epitopes by multiple platform technologies is effective at eliciting high-titer antibody responses against CSP.

We have previously shown that bacteriophage VLPs can induce strong antibody responses in the absence of exogenous adjuvants; co-administration with most standard adjuvant formulations only results in modest boosts in antibody titer ^45^. This is due to the repetitive structure of VLPs, which strongly stimulates B cells ^21, 42, 43^, but also the fact that bacteriophage VLPs encapsidate ssRNA, which can serve as a natural endogenous adjuvant. However, here we showed that combining CIS43 VLPs with delta inulin polysaccharide-based (Advax) adjuvants leads to very robust antibody concentrations that substantially increased protective efficacy in a mouse challenge model. While these high antibody titers were transient, they could be increased by boosting. Importantly, Advax adjuvants have been shown to be safe and efficacious in multiple human vaccine trials ^25^ and recently the FDA approved an IND application for an enhanced seasonal influenza vaccine containing an Advax-CpG formulation.

While the monoclonal-like response conferred by the single epitope-displaying CIS43 VLPs did not confer sterilizing immunity against malaria in a mouse challenge model, the degree of protection elicited by this vaccine highlights the importance of this particular epitope for future clinical applications. Further development of CIS43 VLPs, in combination with vaccines that target other liver- or blood-stage malarial antigens, could increase the breadth and potency of protection.

## Materials and Methods

### Production and characterization of CIS43 epitope displaying bacteriophage Qβ VLPs

Qβ bacteriophage VLPs were produced in *Escherichia coli* (*E. coli*) using methods previously described for the production of bacteriophage PP7 VLPs ^47^. With the exception of the VLPs used in preliminary mouse immunogenicity studies, all CIS43 VLPs and wild-type (WT) Qβ VLPs were depleted of endotoxin by three rounds of phase separation using Triton X-114 (Sigma-Aldrich), as described in reference ^48^. The fifteen amino acid CIS43 epitope peptide was synthesized (GenScript) and modified to contain a C-terminal *gly-gly-gly-cys* linker sequence (NPDPNANPNVDPNAN*GGGC*). The peptide was conjugated directly to surface lysines on Qβ bacteriophage VLPs using the bidirectional crosslinker succinimidyl 6-[(beta-maleimidopropionamido) hexanoate] (SMPH; Thermo Fisher Scientific) as previously described ^47^. The efficiency of conjugation was assessed by gel electrophoresis using a 10% SDS denaturing polyacrylamide gel. Peptide conjugation results in a mobility shift of the Qß coat protein due to an increase in molecular weight. The percentage of coat protein with zero, one, two, or more attached peptides was determined using ImageJ software and used to calculate average peptide density per VLP. Presence of the CIS43 peptide on CIS43 VLPs was confirmed by ELISA. Briefly, 250 ng of VLPs were used to coat wells of an ELISA plate. Wells were probed with dilutions of mAb CIS43 (generously provided by Robert Seder, NIH Vaccine Research Center), followed by a 1:4000 dilution of horseradish peroxidase (HRP) labeled goat anti-human IgG (Jackson Immunoresearch). The reaction was developed using 3,3′,5,5′-tetramethylbenzidine (TMB) substrate (Thermo Fisher Scientific) and stopped using 1% HCl. Reactivity of the CIS43 mAb for the CIS43 VLPs was determined by measuring optical density at 450□nm (OD_450_) using an AccuSkan platereader (Fisher Scientific).

### Ethics Statement for animal studies

All animal research complied with the Institutional Animal Care and Use Committee of the University of New Mexico School of Medicine (Approved protocol #: 19-200870-HSC), Johns Hopkins University (Approved protocol permit #: MO18H419). AlphaGenesis Inc. adheres to the NIH Office of Laboratory Animal Welfare standards (OLAW welfare assurance #A3645-01) and all guidelines of AGI Ethics and Compliance Program were followed.

### Mouse Immunization Studies

Groups of four-week old female Balb/c mice (obtained from the Jackson Laboratory) were immunized intramuscularly with 5µg of CIS43 VLPs or control (unmodified) Qß VLPs. Mice were typically boosted twice after the initial prime, at 3- or 4-week intervals. Some vaccinations were performed using proprietary adjuvants generously provided by Vaxine Ptd Ltd. In these experiments, mice were intramuscularly immunized with 5µg VLPs in combination with 20 µl the following: Advax (50 mg/mL), Advax-2 (50 mg/mL), Advax-3 (5 mg/mL), Advax-4 (50mg/mL), Advax-5 (50mg/mL), and CpG55.2 (0.5 mg/mL) - a synthetic Class B CpG oligonucleotide adjuvant that is dually active on both mouse and human TLR9 (unpublished data). An additional group of mice received 5 µg of unadjuvanted CIS43 VLPs. In these experiments, mice were boosted four weeks after the prime and selected groups received a second boost at 3 months. In each mouse experiment, serum samples were collected prior to each boost and, in some cases, at additional later timepoints following the final boost.

### Cynomolgus Monkey Immunization Studies

Groups (n=3/group) of male and female cynomolgus monkeys (*Macaca fascicularis*) of varying ages (9.9 to 17 years) and body weights (4.00 kg to 13.30 kg) were immunized with 100 µg of unadjuvanted CIS43 VLPs, or the same dose in combination with 0.5 mg of Advax-3. One month after the prime, animals were boosted with 50 µg of VLPs with or without Advax-3. A negative control group received similar doses of unmodified WT Qβ VLPs. Serum was collected at the initial immunization and at two-week intervals thereafter.

### Quantitating antibody responses

Serum antibodies against full-length CSP were detected by ELISA using recombinant CSP expressed in *Pseudomonas fluorescens* ^49^ (and generously provided by Gabriel Gutierrez at Leidos, Inc.) as the coating antigen. Immulon 2 plates (Thermo Scientific) were coated with 250 ng of CSP in 50 µl PBS and incubated at 4°C overnight. Following incubation, wells were blocked with PBS-0.5% milk for 2 hours at room temperature. Sera isolated from immunized animals were serially diluted in PBS-0.5% milk and applied to wells and incubated at room temperature for 2.5 hours. Dilution of 1:4000 of HRP-labeled goat anti-mouse IgG (or, for macaque sera, a 1:4000 dilution of HRP-labeled goat anti-human IgG) was used to detect reactivity to the target antigen. Reactivity was determined using TMB substrate as described above. End-point dilution titer was defined as the greatest sera dilution that yielded an OD_450_ >2-fold over background. For mouse sera, anti-CSP antibody concentrations were also quantitated by ELISA by generating a standard curve using known concentrations of the anti-CSP mouse mAb 2A10 ^7, 30, 48^. For isotype analysis, mouse serum samples were diluted 1:10000, and isotype specific responses were determined using HRP-labeled rat anti-mouse IgG1 or IgG2a (Zymed, diluted 1:4000).

For peptide ELISAs, Immulon 2 plates were coated with 500 ng streptavidin (Invitrogen) for 2 hours at 37°C. Following washing, SMPH was added to wells at 1 μg/well and incubated for 1 hour at room temperature. Specific peptides were added to the wells at 1 μg/well and incubated overnight at 4°C. Wells were then incubated with dilutions of mouse sera and binding was detected as described above.

### Measurement of antibody avidity

The avidity index (AI) of anti-CSP antibodies was evaluated using an ELISA-based chaotrope avidity assay ^52^. This protocol followed the standard ELISA (described above), except that following serum incubation wells were treated with 6M urea for 10 minutes. Serum dilutions corresponding to comparable OD value for each group were chosen to control for differences in antibody concentrations. The AI was calculated as the ratio of the ELISA absorbance value of 6M urea-treated wells (A_6M UREA_) to control wells incubated in water (A_c_); AI = A_6M UREA_/A_c_. Multiple dilutions of sera were analyzed and all samples were tested in duplicate.

### Mouse Pb-PfCSP-Luc sporozoite challenge

Challenge studies were performed using female 6-8 week old C57BL/6 mice. Mice (typically n=7/group) were immunized intramuscularly with 5µg of CIS43 VLPs with or without adjuvant three times at three-week intervals. Separate control groups were immunized with unmodified WT Qß VLPs, PBS, or adjuvant alone. Each immunogen was blinded to minimize the potential for bias in animal handling during the challenge portion of the study. Serum was collected following the third immunization.

Mice were challenged using transgenic *P. berghei* sporozoites engineered to express luciferase and full-length *P. falciparum* CSP in place of *P. berghei* CSP (denoted as *Pb-Pf*CSP*-Luc*) ^53^. Mice were challenged directly using sporozoites or by using infected mosquitos. For the sporozoite challenge, *Pb-Pf*CSP*-Luc* sporozoites were freshly harvested from female *Anopheles stephensi* salivary glands. 1000 sporozoites in 200 µl HBSS/2% FCS were intravenously (i.v.) injected into immunized and naïve mice. 42 hours post challenge, mice were intraperitoneally injected with 100 µl D-luciferin (30mg/ml) and anesthetized. Liver luminescence was assessed by IVIS Spectrum Imaging System (Perkin Elmer). For mosquito challenges, *A. stephensi* mosquitos were infected by blood feeding on *Pb-Pf*CSP*-Luc* infected mice. Prior to challenge, mice were anesthetized with 2% Avertin, and then exposed to 5 mosquitos for a blood meal. Following feeding, mosquitos positive for a blood meal were counted. Liver luminescence was assessed 42 hours post challenge, as described above. Five days later, blood smears were evaluated by Giemsa staining for parasitemia.

### Passive transfer study

Cynomolgus monkey sera was pooled within each group, heat inactivated for 30 minutes at 56°C, and filtered through a 0.45 micron filter to remove aggregates. 500 µl of sera (or PBS as a control) was then passively transferred into each mouse (n=4-5 mice/group) via i.v. tail injection. Two hours following serum transfer, mice were challenged by *A. stephensi* mosquito bite, as described above. Liver luminescence was evaluated 42 hours post challenge, and parasitemia was evaluated by blood smears 4 days later, as described above.

### Statistics

All statistical analyses of data were performed using GraphPad Prism 8.

## Data Availability

The datasets used and/or analyzed in the current study are available from the corresponding author upon reasonable request.

## Acknowledgements

This study was supported, in part, by a generous contribution to the UNM Foundation in honor of Jeffrey Michael Gorvetzian in support of biomedical research excellence at the University of New Mexico School of Medicine. FZ, HJ and AE thank the Bloomberg Philanthropies for continued support. Development of Advax adjuvants was supported by funding from National Institute of Allergy and Infectious Diseases of the National Institutes of Health under Contracts HHS-N272201400053C, HHS-N272200800039C and U01-AI061142.

## Author Contributions

L.J., F.Z., N.P. and B.C. designed the study and drafted the manuscript. L.J. engineered the VLPs, performed the mouse experiments at the University of New Mexico, performed the analysis of immune responses, and analyzed all data. H.J., A.E. and F.Z. performed the mouse challenge experiments and statistically analyzed the data resulting from those experiments. N.P. supplied adjuvants and coordinated the macaque immunizations.

## Supporting information

**Supplementary Figure 1.**
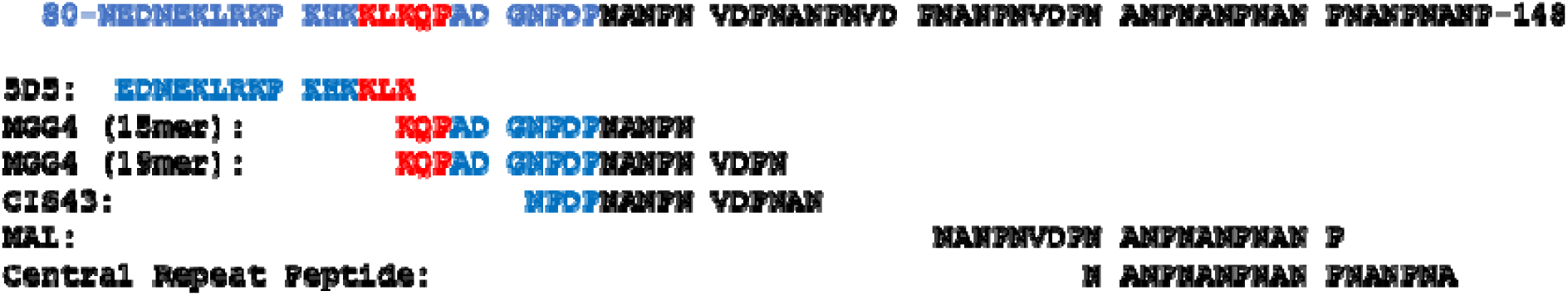
Sequence of the CSP junctional domain and the peptides used in this study. Colors indicate the N-terminal domain (blue), R1 cleavage site (red), and repeat domain (black). All peptides were also synthesized to contain a C-terminal *-GGGC* sequence to facilitate crosslinking.

**Supplementary Figure 2.**
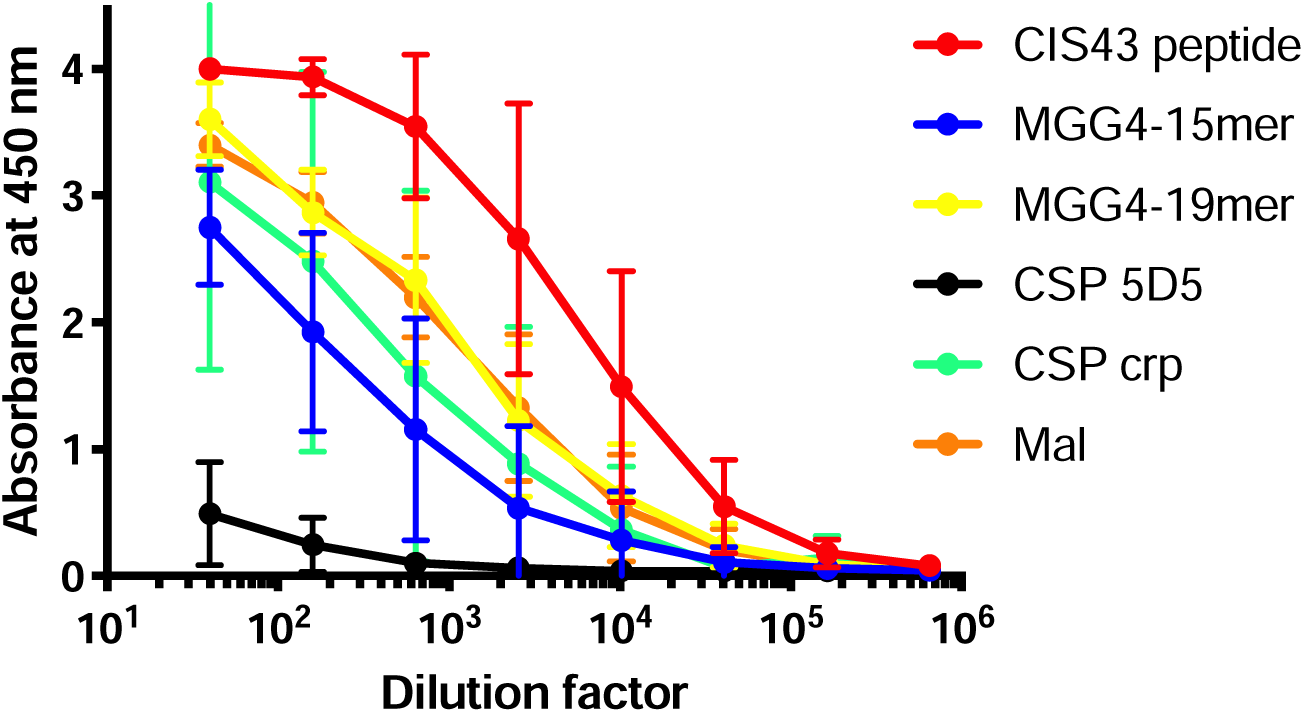
Sera from CIS43 VLP-immunized mice binds promiscuously to diverse CSP peptides. Sera from CIS43 VLP immunized Balb/c mice were tested for binding to different CSP peptides (shown in Supplementary Figure 1) by peptide ELISA. Data represents the mean values from five mice immunized twice with unadjuvanted CIS43 VLPs. Error bars show standard deviation (SD).

**Supplementary Figure 3.**
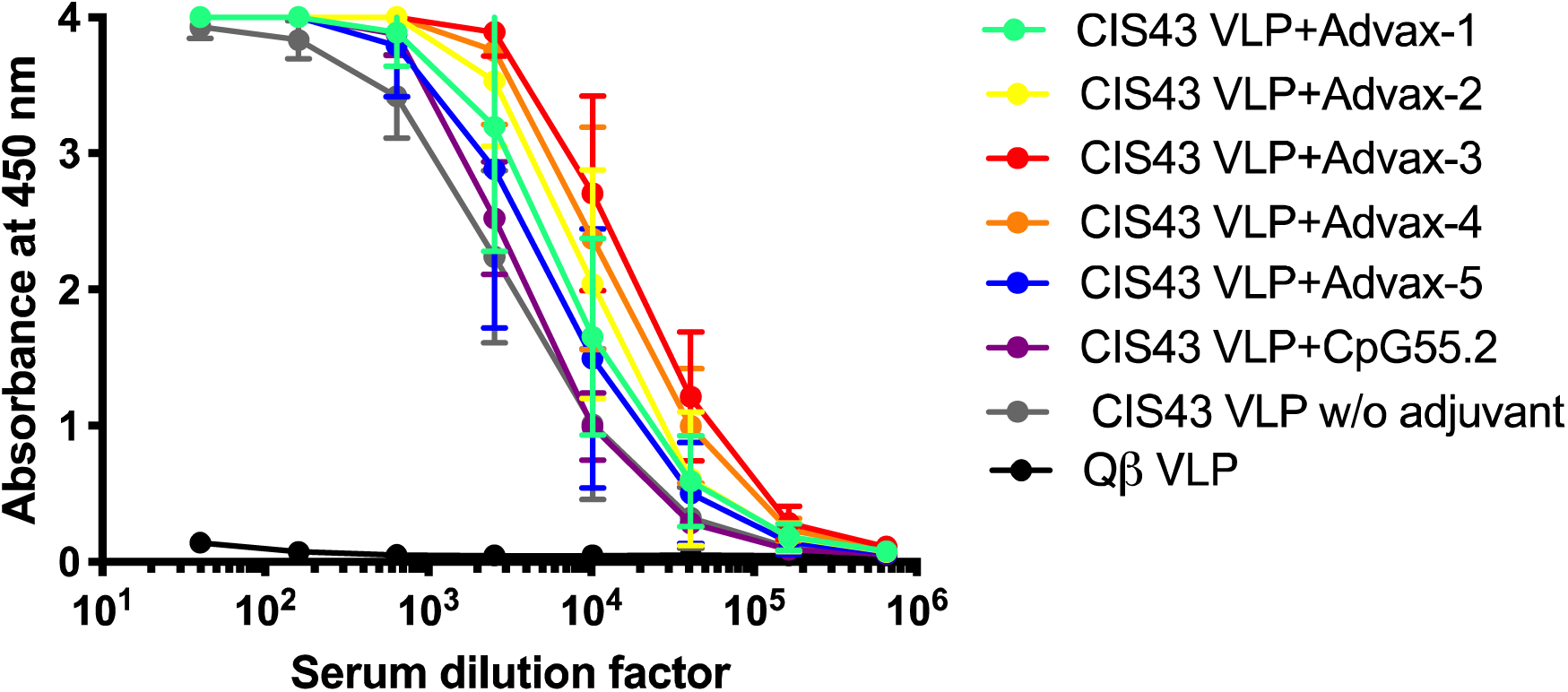
Advax adjuvants enhance anti-CSP IgG response to CIS43 VLP immunization. Groups of Balb/c mice (n = 5) were immunized with two doses of CIS43 VLPs adjuvated with different Advax adjuvant formulations or CpG oligonucleotides. Unadjuvanted CIS43 VLPs and WT Qß VLPs were used as controls. Serum was collected three weeks following the second immunization.

**Supplementary Figure 4.**
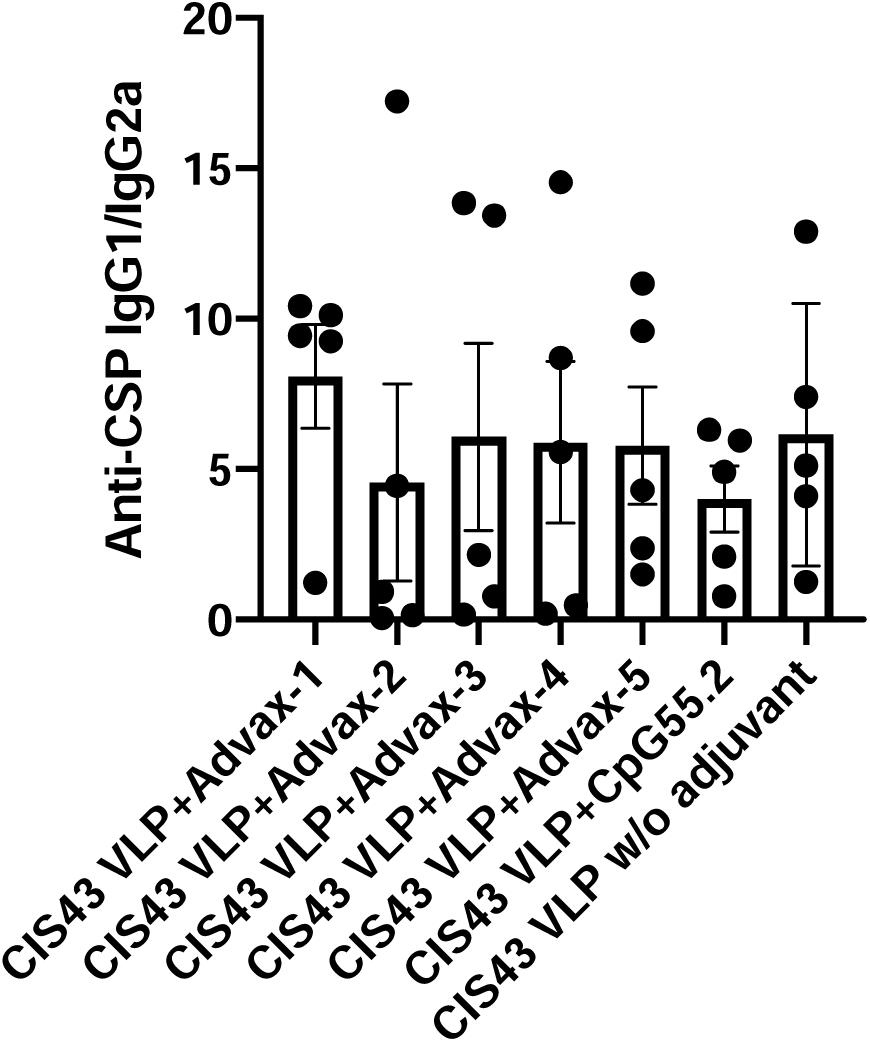
Ratio of detected IgG1 to IgG2A in serum of mice immunized with adjuvanted CIS43 VLPs. No statistical differences were detected in anti-CSP IgG isotypes in mice immunized with CIS43 VLPs adjuvanted with Advax-1, Advax-2, Advax-3, Advax-4, Advax-5, or CpG55.2 compared to those immunized with unadjuvanted CIS43 VLPs.

**Supplementary Table 1.**
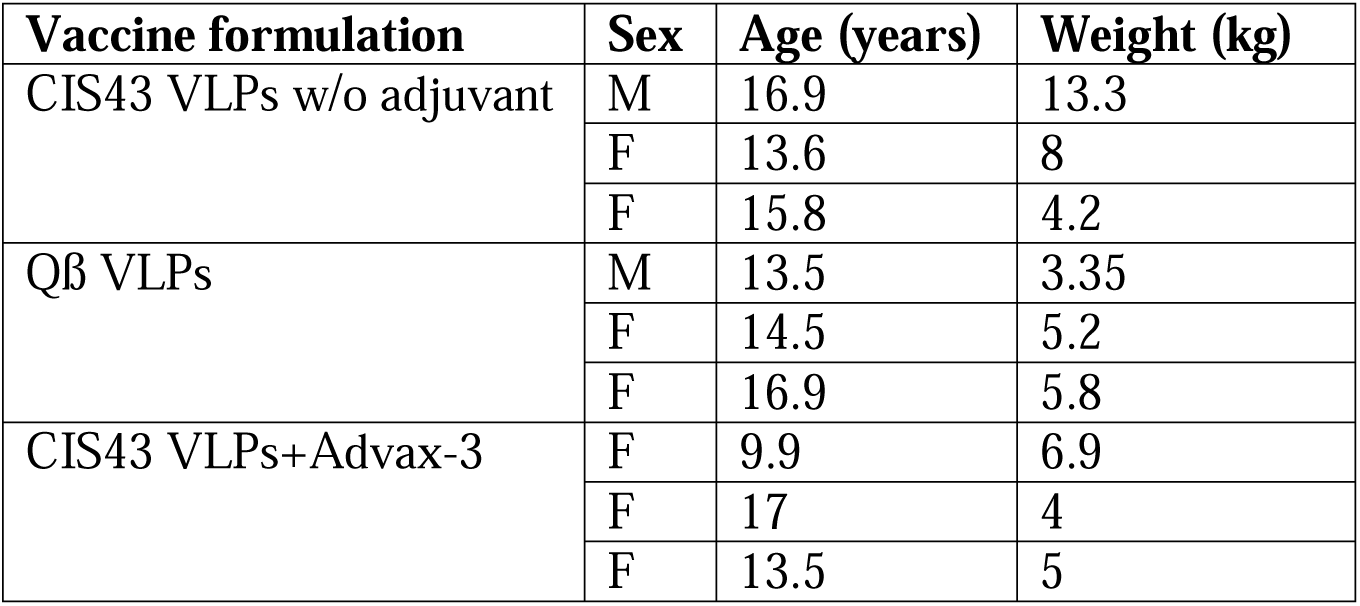
Demographic information of cynomolgus monkeys in the three vaccine groups.

